# Physiological and morphological correlates of extreme acid tolerance in larvae of the acidophilic amphibian *Litoria cooloolensis* (Anura: Pelodryadidae)

**DOI:** 10.1101/2019.12.15.872259

**Authors:** Edward A. Meyer, Craig E. Franklin, Rebecca L. Cramp

## Abstract

In the coastal sandy lowlands, or wallum, of east Australia, a number of anuran species show remarkable tolerance to dilute waters of low pH, including the Cooloola Sedgefrog *Litoria cooloolensis*, larvae of which inhabit water bodies as acidic as pH 3.5. To investigate the physiological and morphological underpinnings of larval acid tolerance in *L. cooloolensis*, we compared survivorship, Na^+^ balance, uptake and efflux rates, and gill and skin morphology in *L. cooloolensis* larvae reared in circum-neutral (pH 6.5) and pH 3.5 water. We hypothesised that larvae reared at pH 6.5 and acutely transferred to pH 3.5 would show an initial loss of ionic homeostasis but with chronic exposure would restore sodium balance, reducing sodium efflux and increasing sodium uptake from the environment. We found that net Na^+^ flux rates were not significantly different from zero in larvae reared at pH 3.5 or in acid-naïve animals maintained in pH 6.5 water. Animals reared at pH 6.5 and acutely exposed to pH 3.5 did, however, exhibit a net loss of Na^+^, due to significant inhibition of Na^+^ uptake. *L. cooloolensis* larvae reared at pH 3.5 were able to maintain Na^+^ balance at pH 3.5 and did not exhibit inhibition of Na^+^ uptake at this pH. Detailed investigation of Na^+^ transport kinetics and the morphology of the gills and integument of *L. cooloolensis* larvae suggests tolerance of *L. cooloolensis* larvae to low pH may be attributed to a high capacity for branchial Na^+^ uptake, increased tight junction length and elevated mucus production in the gills and integument. These factors promote resistance to acid damage and disruption of ionic homeostasis which would otherwise result in the death of amphibian larvae exposed to waters pH 4.0 and less.

## Introduction

For fish and aquatic amphibian larvae survival in acidic waters presents a significant challenge with waters pH 5.0 and less inhibiting active Na^+^ uptake, damaging gill epithelia, and increasing diffusive efflux of electrolytes across the gills and integument (Ferreira and Hill, 1982; Freda and Dunson, 1984; Freda et al., 1991; McDonald, 1983; Meyer et al., 2010). Damage to the gills of fish and amphibian larvae exposed to low pH includes lifting of the branchial epithelium, dilation of paracellular spaces and opening/shortening of tight junctions between gill pavement cells (Daye and Garside, 1976; Meyer et al., 2010; Rosseland and Staurnes, 1994). The exposure of fish and amphibian larvae to low pH water can also result in significant damage to the integument (e.g. tight junction damage and cell swelling and sloughing and localised necrosis) and in amphibian larvae, it is the integument which appears most sensitive to acid damage (Daye and Garside, 1976; Meyer et al., 2010). With the structural integrity of epithelia compromised, transepithelial resistance to ionic efflux is greatly reduced. The resultant net loss of plasma electrolytes (in particular Na^+^) can lead to substantial ionic and osmotic disturbances, with the loss of 50% of body Na^+^ resulting in death (Freda and Dunson, 1984; Freda et al., 1991; Laurent and Perry, 1991; McDonald, 1983; Meyer et al., 2010; Milligan and Wood, 1982; Rosseland and Staurnes, 1994). Problems with electrolyte loss appear to be exacerbated in soft (Ca^2+^-poor) waters where reduced availability of Ca^2+^ promotes diffusive ionic efflux (Freda and Dunson, 1984; McDonald, 1983).

Despite the damaging effects of low pH on epithelial structure and function, a number of fish and amphibian species tolerate acidic conditions remarkably well (Gonzalez et al., 2002). In the coastal sandy lowlands of east Australia, surface waters are naturally soft and acidic with low concentrations of key electrolytes such as Na+ and Cl-(Bensink and Burton, 1975; Hines and Meyer, 2011). With relatively soft waters of such low pH, the maintenance of ionic homeostasis and survival in these areas presents a significant challenge to aquatically respiring animals. Yet despite this, frog species endemic to these regions, breed successfully in soft waters of ≤ pH4.0 (Hines and Meyer, 2011; Ingram and Corben, 1975). Like animals from other naturally acidic environments, e.g., fish from the ion-poor acidic Rio Negro (Gonzalez et al., 1997), larvae of these frogs appear be able to resist the damaging actions of acidic soft waters.

The ability of acidophilic fish to maintain ionic homeostasis and survive at low pHs (in particular fish species native to the Rio Negro) has attracted considerable interest (Gonzalez et al., 1997; Gonzalez and Preest, 1999; Gonzalez and Wilson, 2001; Gonzalez et al., 2002; Wilson et al., 1999; Wood et al., 2003). In these animals, acid tolerance is conferred by physiological and/or morphological mechanisms that allow animals to resist, or compensate for, the disruption of ionic homeostasis and associated damage to the gills (Gonzalez et al., 1997; Gonzalez and Dunson, 1989b; Gonzalez and Preest, 1999). In acid tolerant fish, resistance to low pH has been linked to a high branchial affinity for Na^+^ and/or increased rate of Na^+^ uptake, allowing for the maintenance of Na^+^ uptake at low pHs (Gonzalez et al., 1997; Gonzalez and Preest, 1999). Tolerance of fish to acid exposure has also been linked to control of paracellular ionic permeability, with greater branchial tight junction length promoting resistance to increased ionic efflux at low pH (McDonald et al., 1991). Studies of acid tolerant fish also suggest that control of ionic efflux at low pH may be linked to a high branchial affinity for Ca^2+^, with calcium playing an important role in maintaining transepithelial resistance (see, e.g., Gonzalez et al., 1997; Gonzalez and Preest, 1999). In addition, resistance to acid damage and control branchial ionic efflux at low pH has also been tied to increased mucus production at the gills with increased mucus secretion likely to protect branchial epithelia and tight junctions from acid damage (Jagoe and Haines, 1983; McDonald, 1983; McDonald et al., 1991).

By comparison, very little has been published on acid tolerance in amphibian species, with studies on anuran larvae dealing mainly with effects of short-term (acute) acid exposure on ion regulation in acid-sensitive species (Freda and Dunson, 1984; McDonald et al., 1984; Meyer et al., 2010). These studies tell us little about the mechanisms underpinning extreme soft water acid tolerance in acidophilic frog species. In this study we investigated physiological and morphological mechanisms for extreme acid tolerance in the Cooloola sedgefrog (*Litoria cooloolensis*), one of a number of ‘acid’ frog species endemic to the ‘wallum’ of coastal eastern Australia, an area of sand plains and dunes with swamps and lakes contain dilute (soft) water as acidic as pH 3.0 (Hines and Meyer, 2011). We hypothesised that relative to the closely-related but acid sensitive species, *Litoria fallax* (Meyer et al., 2010), larvae of the acid tolerant *L. cooloolensis* would exhibit distinct physiological and morphological traits that permit the maintenance of ionic homeostasis at low pH.

## Materials and Methods

### Experimental animals

Larvae used in laboratory experiments were reared from spawn collected from four pairs of amplectant *L. cooloolensis* captured at Lake Poona, Cooloola National Park and Brown Lake, North Stradbroke Island (Queensland, Australia). All amplectant animals were released at the point of capture following the collection of spawn. Water samples from capture sites were collected and analysed by ICPOES / ICPMS (School of Agriculture and Food Sciences, The University of Queensland; Table 1).

**Table 1.** Water chemistry data for *Litoria cooloolensis* egg collection sites. Data are presented as the range of values measured (min – max).

| Water Attribute | Range |
| --- | --- |
| pH | 4.4 - 4.8 |
| Ca<sup>2+</sup> (μmol L<sup>-1</sup>) | 15 - 33 |
| Mg<sup>2+</sup> (μmol L<sup>-1</sup>) | 132 - 210 |
| Na<sup>+</sup> (μmol L<sup>-1</sup>) | 74 - 195 |
| K<sup>+</sup> (μmol L<sup>-1</sup>) | 30 - 40 |
| Cl<sup>-</sup> (μmol L<sup>-1</sup>) | 485 - 860 |

Fifteen to twenty-five embryos at Gosner (1960) stages 9-10 from each clutch were transferred to individual four litre tubs containing two litres of artificial soft water (ASW; distilled water plus (in mmol L^-1^) 0.04 CaCl_2_.2H_2_O, 0.04 MgSO_4_.7H_2_O, 0.12 NaCl, 0.05 NaOH, 0.02 KCl) with the pH adjusted to 6.5 or 3.5 through the addition of dilute (0.1 M) sulphuric acid. Larvae reared in pH 6.5 water and 3.5 water are hereafter referred to as ‘acid naïve’ and ‘acid acclimated’, respectively. Larvae were reared under these conditions until Gosner stage 25-30 (Gosner, 1960). During this time, water pH was checked twice daily with a temperature compensated pH meter (Hannah Instruments, Melbourne, VIC, Australia) and, where necessary, adjusted with 0.1 M sulphuric acid. During the course of experiments, pH did not drift by more than 0.2 units. Larvae were fed a mixture of frozen boiled endive and lettuce *ad libitum*. Leftover food and faeces were removed from containers daily and a third of the water in containers replaced every other day. Larvae were reared at room temperature (23 - 25 °C) with a 12 h light: 12 h dark photoperiod. Larvae used in experiments were matched for size (mean wet mass ± s.e.m. = 0.147 ± 0.007 g [N = 105]). All animals used in experiments were starved for three days prior to experimentation.

### Effects of acute and chronic exposure to low pH on body Na^+^ and water contents

To assess the sensitivity of larvae to acute acid exposure, twenty acid naïve larvae were removed from their holding tubs and placed in separate 250 mL glass beakers filled with fresh pH 6.5 ASW. Ten acid-acclimated larvae were also removed from holding tanks and placed into individual beakers containing fresh pH 3.5 ASW. After 24 hours, the pH of water in half of the beakers containing naïve larvae was acutely lowered to 3.5 ± 0.05 through the addition 0.1 M sulfuric acid. The pH of the water of remaining naïve larvae and the acid acclimated larvae was not changed. Every half hour thereafter, larval survival was assessed by prodding the larvae gently with a plastic pipette to determine whether they exhibited the natural startle/escape response. Unresponsive larvae were removed from beakers, blotted dry, weighed and analysed for tissue Na^+^ content (below). After 24 hours, all surviving larvae were sacrificed with ethyl 3-aminobenzoate methanesulfonate (MS 222; diluted 1:2000), blotted dry, weighed and analysed for tissue Na^+^ content. For the determination of body Na^+^ content, larvae were placed inside pre-weighed, acid-washed glass test tubes and dried to a constant mass at 95 ^o^C and re-weighed. Tissue water content (%) was determined by subtracting the larval dry mass from the wet mass. Dried larvae were then dissolved in 3 mL of concentrated nitric acid. After three days, 1 mL of digest was placed into an acid-washed plastic tube and eight mL of distilled water was then added to the digest in test tube and the Na^+^ concentration of diluted samples determined via plasma-coupled atomic emission spectroscopy (Queensland Health Laboratories, Nathan, QLD, Australia).

### Na^+^ transport measurements

For acid naïve larvae, rates of Na^+^ uptake (J_in)_, Na^+^ efflux (J_out_) and net Na^+^ flux (J_net_) were measured at both their acclimation pH (6.5) and following acute exposure to pH 3.5 water. Na^+^ fluxes were measured in acid acclimated larvae at their acclimation pH of 3.5 only. Larvae were left overnight in individual 40 mL glass flux chambers connected to a 15 L re-circulating system and supplied with pH 6.5 (acid naïve larvae) or pH 3.5 (acid acclimated larvae) ASW at a rate of 20 mL h^-1^. The following morning, flow to all chambers was stopped and 15 kBq.L^-1^ of ^22^NaCl was added to each chamber; for the measurement of Na^+^ fluxes in acid naïve larvae at pH 3.5, dilute 0.1 M sulfuric acid was added to chambers prior to the addition of ^22^NaCl. After 5 min, an initial 10 mL water sample was withdrawn from each chamber. An hour later a second 10 mL water sample was withdrawn and tadpoles were removed from flux chambers. Larvae were rinsed in unlabelled ASW water and transferred to 7 mL plastic test tubes for direct counting in an LKB 1277 Gamma Master (LKB Wallac, Turku, Finland). Water samples were similarly assayed for ^22^Na-activity. Once counted, water samples were analysed for Na^+^ concentration via ICAEP. Net Na^+^ flux (J_net_), Na^+^ uptake (J_in_), and Na^+^ loss (J_out_), were calculated following Gonzalez et al. (1997):

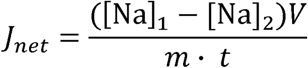

where [Na]_1_ and [Na]_2_ are the Na^+^ concentrations in the flux champers at the beginning and end of the flux period (in mmol L^-1^), V is the flux chamber volume in L, m is the mass of the tadpole in grams and t in the time in hours. Na^+^ influx (J_in_) was calculated using the equation:

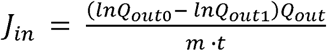

where Q_out0_ and Q_out1_ are the total counts per minute in the flux chambers at the beginning and end of the flux period, respectively; *Q*_out_ is the average amount of Na^+^ in the flux bath during the flux period, m is the mass of the tadpole in grams and t is the time in hours. J_out_ (Na^+^ efflux rate) was calculated as the difference between J_net_ and J_in_.

To investigate the role of Ca^2+^ in controlling ionic efflux at low pH, the influence of external calcium concentration on Na^+^ efflux rates were compared in larvae reared at pH 6.5 and acutely exposed to pH 3.5 water and larvae reared and tested in pH 3.5 water. Using the protocol detailed above, external calcium concentrations were adjusted to approximately twice and ten times the amount of Ca^2+^ in ASW (i.e., with [Ca^2+^] = 0.08 mmol L^-1^ and 0.4 mmol L^-1^) in the flux chambers immediately prior to the reduction in chamber pH and the addition of the radioisotope. Na^+^ efflux rates were calculated as detailed above.

Na^+^ transport kinetics (affinity and maximum uptake rates) were assessed by measuring the dependence of Na^+^ uptake on environmental Na^+^ concentrations in both larvae reared at pH 6.5 and pH 3.5. All measurements were made at pH 6.5. Prior to measurements, larvae were isolated overnight in glass beakers containing 200 mL ASW at pH 6.5. The following morning, tadpoles were transferred to glass beakers with 50 mL radio-labeled pH 6.5 ASW (^22^NaCl: 15 kBq L^-1^) with Na^+^ concentrations ranging from 0.05 mmol L^-1^ through to 3.5 mmol L^-1^ (the concentration of Na^+^ was modified through the omission or addition of NaCl from or to ASW). Five to seven larvae of each species were exposed to each of five different sodium concentrations. After 1 h, tadpoles were removed from beakers and rinsed in unlabeled pH 6.5 water for 1 min each. Thereafter tadpoles were counted using a Packard gamma counter. The transport mechanism’s affinity for Na^+^ (K_m_), and maximum transport capacity, J_max_, were estimated using non-linear regression. The regression model for estimating these parameters was:

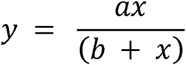

where y = J_in_, *a* = J_max_; *b* = K_m_ and *x* = [Na^+^]_e_ (the concentration of Na^+^ in bath water).

### Morphological Analyses

Gills and integument of *L. cooloolensis* larvae reared from embryos at pH 6.5 and pH 3.5 (Gosner stage 25-29) were examined under light (LM), transmission electron (TEM) and scanning electron microscopes (SEM). Gills and a section of tail were excised from larvae sacrificed with MS222 (1:2000). Tissues for LM were fixed in 10% neutral buffered formaldehyde for 24 h, dehydrated and cleared with ethanol and xylene, respectively, and embedded in paraffin wax. Blocks were sectioned at 6 µm and stained with hematoxylin and eosin. Tissues for TEM were fixed for 2 h in cold 2.5% glutaraldehyde in 0.1M cacodylate buffer. Tissues were post-fixed in osmium tetroxide (1% solution in 0.1 M cacodylate buffer), dehydrated in an ascending ethanol series and embedded in Spurr’s resin. Ultrathin sections were stained as per Daddow (1986) in lead citrate and uranyl acetate, and viewed and photographed with a Hitachi 300 TEM operating at 75 kV. For SEM, tissue samples were fixed for 2 h in 2.5% glutaraldehyde in 0.1M cacodylate buffer, dehydrated in ethanol and critical point dried with liquid CO_2_ (Polaron; ProSciTech, Thuringowa, QLD, Australia). Whole gills, gill arches and tail sections from acid naïve and acid acclimated larvae (N = 3 for each group) were mounted on stubs and coated with gold in a sputter coater (SPI Module; SPI Supplies, West Chester, PA, USA). Gills and tail sections were viewed and photographed with a Jeol JSM 6400 SEM.

The gross morphology of gill tufts and the integument was viewed using SEM and LM. A qualitative assessment, based on SEM and LM examination of tissues, was made of gill tuft proportions (height and thickness), lymphatic/paracellular space, and thickness of the integument. Gill tuft sections were viewed at low power (5000 – 10 000 x magnification) with a Hitachi 300 transmission electron microscope operating at 75 kV. Intercellular junctions were measured from digital TEM images using a Warcom graphics tablet and image analysis software (SigmaScanPro™). The junctional attributes measured were total length of the junctional complex between cells (including both the tight and adherens junctions, tight junction length, minimum width (the distance between adjacent cell membranes within the tight junction) and maximum width (the distance between apposing cell membranes within the adherens junction). Gill tuft pavement cells (between 10 and 15 per animal) and epithelial cells of the integument (between 10 and 15 per animal) were examined using TEM and the abundance of patent mucus secretory vesicles at the apical surface of cells, secretory vesicle cross-sectional area and cell cross-sectional areas were measured (SigmaScanPro^™^).

### Statistical analyses

Mean body Na^+^ and water contents were compared using t-tests. Mean J_net_, J_in_ and J_out_ of acid naïve and acid acclimated larvae at their acclimation pH and in acid naïve larvae acutely exposed to pH 3.5 ASW were compared using one-way ANOVA and Sidak post-hoc tests. Mean J_net_ values were judged significantly different from zero using one sample means t-tests. A Linear mixed-effects model with Tukeys post-hoc tests was used to compare mean J_out_ at different calcium concentrations across acclimation treatments. Where necessary, data were log or arcsine-square root transformed prior to analysis. For the morphological variables, we sought to identify morphological differences between acid naïve and acid acclimated larvae reflecting acclimation with chronic exposure to pH 3.5 soft water. All morphological parameters were compared using MANOVA (α = 0.05). Unless otherwise stated all values reported are means ± s.e.m. Analyses were conducted using GraphPad Prism 8.3.

## Results

### Survivorship, body Na^+^ and water contents

Acid naïve *L. cooloolensis* larvae acutely exposed to pH 3.5 water suffered 20% mortality after 24 hours; however, acid naïve and acid acclimated larvae maintained at their acclimation pH experienced no mortality during this period. The body Na^+^ content of acid-naïve larvae acutely exposed to pH 3.5 water was significantly lower than that of larvae maintained at pH 6.5, indicating a net loss of body Na^+^. Acid-naïve larvae acutely exposed to pH 3.5 that died within 24 h had lost > 60 % of total body Na^+^ (388 ± 20 µmol Na^+^ g^-1^ DBM), while those that survived for 24 h at pH 3.5 lost on average 35 % of their body Na^+^ (620 ± 30 µmol Na^+^ g^-1^ DBM; *t* = 5.54, *df* = 27, *P* < 0.001). The total body Na^+^ content of *L. cooloolensis* larvae reared at pH 3.5 was slightly, but not significantly (∼ 11 %) lower than that of larvae reared at pH 6.5 (862 ± 45 and 969 ± 55 µmol Na^+^ g^-1^ dry body mass [DBM], respectively; *t* = 1.7, *df* = 27, *P* = 0.1). Body water contents did not differ significantly between pH treatments with water comprising approximately 90% of wet body mass of all larvae. The dry mass of gills (as a percentage of dry body mass) was not significantly different between acid naïve and acid acclimated larvae (0.62 ± 0.09 and 0.76 ± 0.06 %, respectively).

### Na^+^ fluxes

For acid naïve larvae tested at pH 6.5 rates of Na^+^ uptake (J_in_) and efflux (J_out_) were roughly equal such that J_net_ was not significantly different from zero (acid naïve: *t* = 1.12, *df* = 5, *P* = 0.32; Fig 1a). This was also the case for acid acclimated larvae tested at pH 3.5 (acid acclimated: *t*= 2.4, *df* = 6, *P* = 0.06). Mean Na^+^ influx rates for acid acclimated larvae at pH 3.5 were slightly (20%), but not significantly, higher than those of acid naïve larvae at pH 6.5 (J_in_: *t* = 1.77, *df* = 15, *P* = 0.18). However, rates of Na^+^ efflux were ∼30 % higher in acid acclimated larvae (J_out_: *t* = 3.28, *df* = 15, *P* = 0.01). Relative to larvae at pH 6.5, acute exposure of acid naïve larvae to pH 3.5 water resulted in a 50% decrease in Na^+^ uptake rate (*t* = 6.23, *df* =15, *P* < 0.001) but had no effect on Na^+^ efflux rates (*t* = 0.92, *df* = 18, *P* = 0.62). This reduction in uptake resulted in a significant net loss of Na^+^ (*t* = 3.77, *df* = 6, *P* = 0.009).

**Figure 1.**
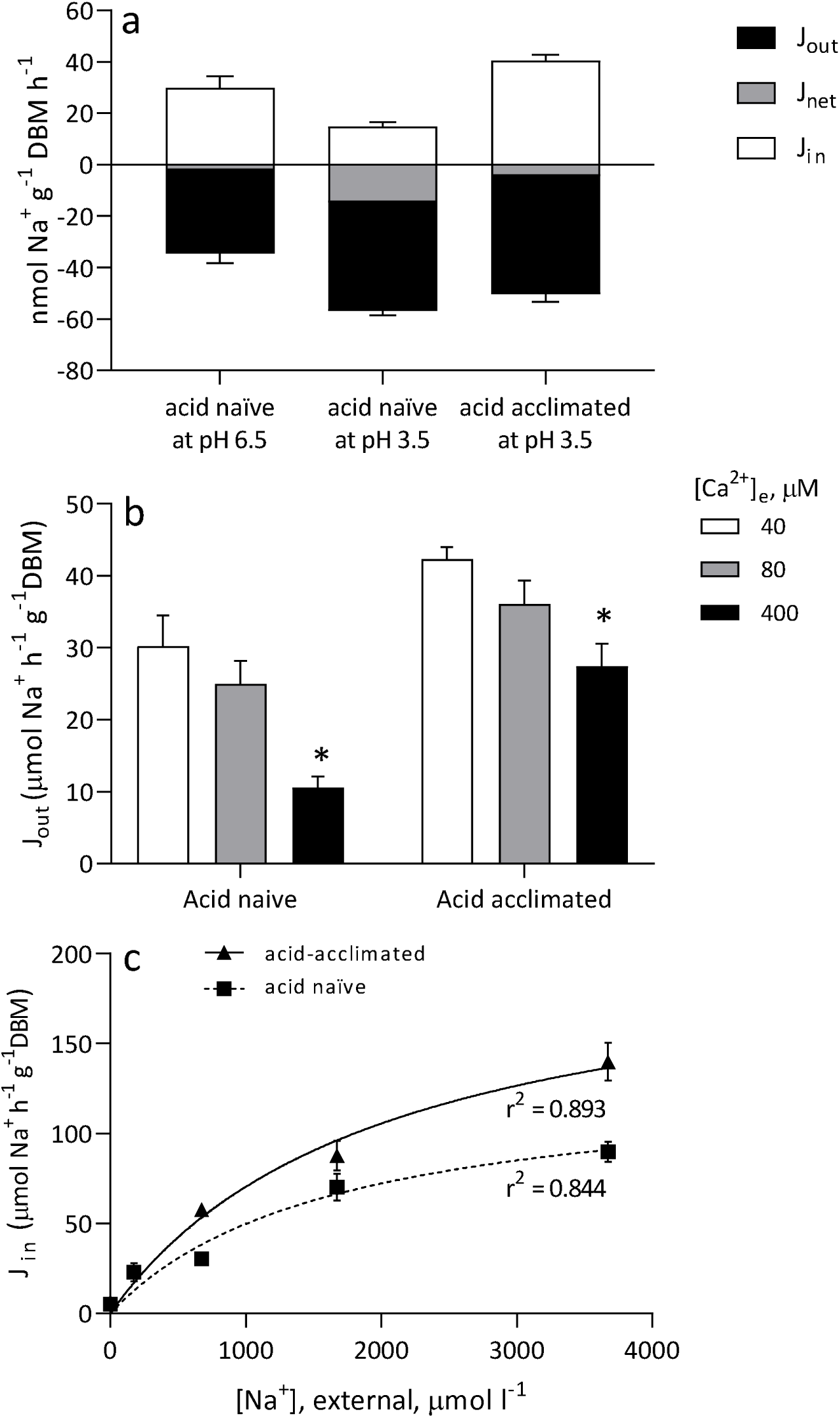
(a) Na^+^ flux parameters (influx - J_in_; efflux – J_out_; net flux – J_net_) in *L. cooloolensis* larvae tested in artificial soft water (ASW) at their acclimation pH (acid naïve larvae = pH 6.5; acid acclimated larvae = pH 3.5) and for acid naïve larvae, acutely at pH 3.5. J_in_ data are presented as positive values and J_out_ data are presented as negative values for the purposes of visual representation. N = 6-7 animals per treatment group. (b) The effect of rearing pH and environmental Ca^2+^ level on Na^+^ efflux rates in *L. cooloolensis* larvae tested at pH 3.5. * indicates that within each acclimation treatment, value is significantly different (P < 0.05) from the rate in ‘control’ (40 μM Ca^2+^) ASW. N = 6 - 7 per treatment. (c) The dependence of Na^+^ influx at pH 6.5 on environmental Na^+^ concentration. Both acclimation treatments were tested at pH 6.5. Lines represent the output of non-linear regression analyses. N = 5 - 7 per [Na^+^] per species. All data are presented as means ± s.e.m.

pH acclimation group (*F*_*(1,17)*_ = 39.39, *P* < 0.001) and water calcium concentrations had significant main effects on Na^+^ efflux rates in *L. cooloolensis* larvae (*F*_*(2,17)*_ = 19.15, *P* < 0.001); there was also a significant interaction between the two factors (*F*_*(2,14)*_ = 4.72, *P* = 0.027). In both acid naïve and acid acclimated larvae, Na^+^ efflux rates at pH 3.5 were unaffected by increasing water calcium levels to 80 μM, but were significantly reduced by 66 and 35% respectively (compared to 40μM Ca^2+^), when [Ca^2+^]_e_ was increased to 400 μM (acid naïve: *q* = 9.4, *df* = 31, *P* < 0.001; acid acclimated: *q* = 3.5, *df* = 31, *P* = 0.048; Fig 1b).

In all larvae, Na^+^ uptake increased with increasing water Na^+^ content ([Na^+^]_e_). J_max_ of acid acclimated larvae at pH 6.5 (210.6 ± 26.32 μmol Na^+^ g^-1^ DBM h^-1^) was significantly greater than that of *L. cooloolensis* 6.5 larvae at pH 6.5 (131.8 ± 18.72 μmol Na^+^ g^-1^ DBM h^-1^; *F*_(1,62)_ = 4.29, *P* = 0.042). However, the K_m_ of acid acclimated larvae (1648 ± 543.6 µmol l^-1^) was not significantly different from that of acid naïve *L. cooloolensis* larvae (1990 ± 529.1 µmol l^-1^; *F*_(1,62)_ = 0.17, *P* = 0.68; Fig 1c).

### Gill and Integument morphology

The gills of both acid acclimated and acid naïve *L. cooloolensis* larvae were structurally very similar. The gills comprised four arches with thin-walled, vascularised tufts projecting ventrally from each arch and filter plates situated dorsally (Fig. 2a). The epithelium of gill tufts comprised a single layer of flattened pavement cells and mitochondrion-rich (MR) cells held together by tight and adherens junctions and desmosomes (Figs. 2b-f). Pavement cells, the dominant cell type, were generally flat. Visible within the cytoplasm of these cells were mitochondria and numerous mucus secretory vesicles. Mucus released from these vesicles adhered to the surface of gill tufts. Dark-staining MR cells, which were few in number, were packed with mitochondria and contained few mucus secretory vesicles. The thickness of the gill tuft epithelium and gill tuft proportions (thickness and relative length) were similar in both acid naïve and acid acclimated larvae.

**Figure 2.**
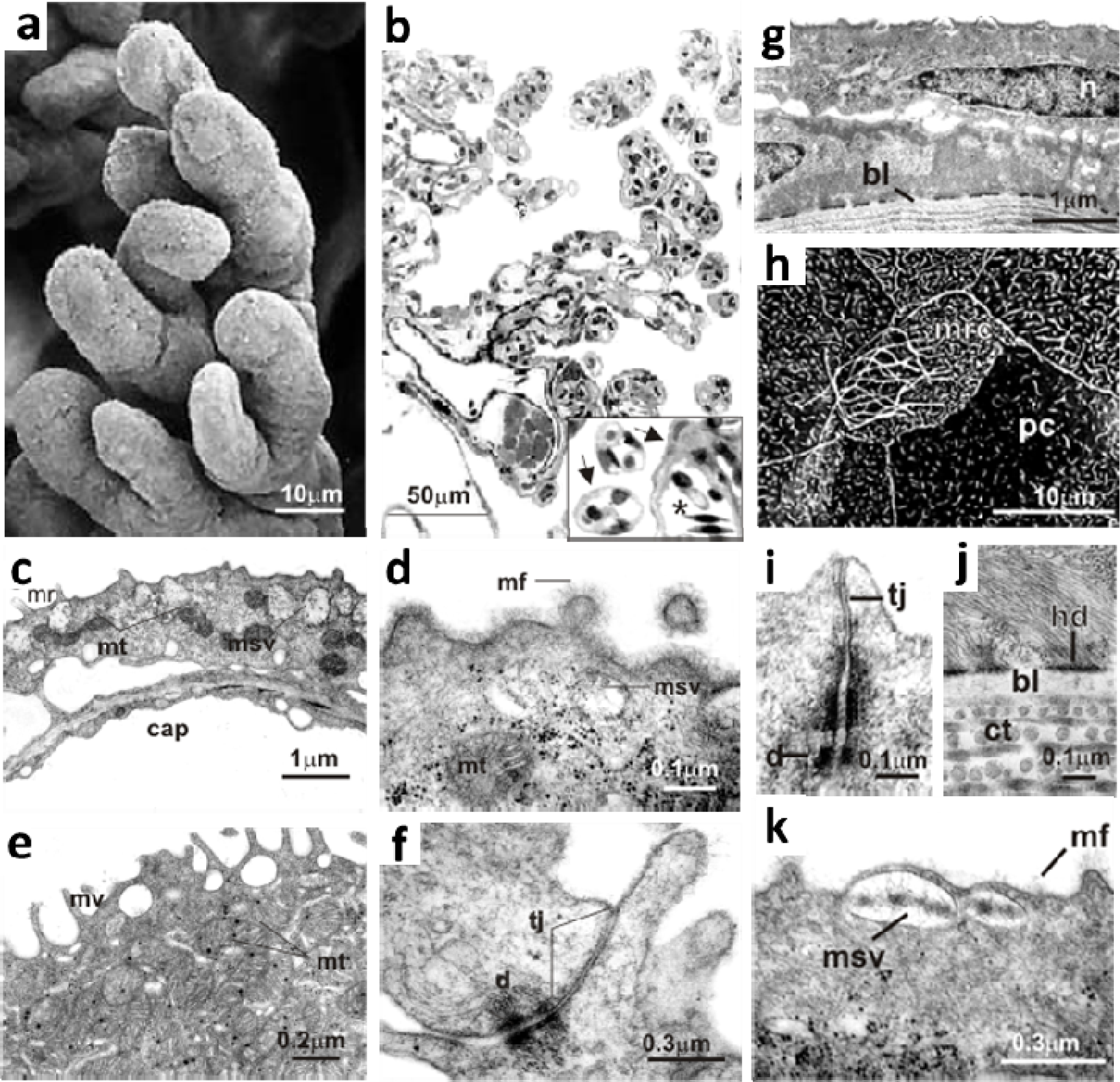
Branchial (a-f) and integument (g-k) morphology and ultrastructure. (a) Gill tufts from an acid naïve larva (b) Cross section of gill of *L. cooloolensis* larva in transverse view; (b) Gill tuft in transverse and oblique view; (c) Gill tuft pavement cell with mucus secretory vesicles and mitochondria; (d) Branchial junctional complex between adjacent pavement cells; (e) Apical surface of gill pavement cell coated with mucus; (f) Apical surface and cytoplasm of mitochondria-rich (MR) cell; (g) Transverse section through the integument of acid naïve larva; (h) Apical surface of squamous epithelial cell of the integument; (i) Attachment of basal cell layer of integument to basal lamina; (j) Junctional complex between adjacent epithelial cells in the integument; (k) SEM micrograph showing apical surface of integument. Abbreviations: bl = basal lamina; cap = capillary; cc = ciliated cell; ct = connective tissue; d = desmosome; ec = squamous epithelial cell of outer integument; fp = filter plates; ga = gill arch; gt = gill tuft; hd = hemi-desmosome; mf = mucus fuzz; mr = microridge; msv = mucus secretory vesicle; mt = mitochondria; n = nucleus; pc = pavement cell; rbc = red blood cell; tj = tight junction; v = velum.

The epidermis of *L. cooloolensis* larvae consisted of a double layer of flattened squamous epithelial cells above a basal lamina (Fig. 2g). The apical membrane of cells in the outer epidermis was amplified with microridges (pavement cells) and microvilli and, in some cases, cilia as well (Fig. 2h). Epidermal cells were held together by desmosomes and tight junctions (Fig. 2i). Cells of the basal layer were attached to the basal lamina by half-desmosomes (Fig. 2j). Cells of the outer layer contained numerous mucus secretory vesicles (Fig. 2k). Electron-dense mucus released from these vesicles adhered to the apical surface of cells.

Branchial tight junction length and total branchial junction lengths were significantly greater (35% and 20% greater, respectively) in acid acclimated larvae relative to acid naïve larvae (Table 2). Minimum width and maximum width were also greater (20 %) in acid acclimated larvae. Similarly, maximum junctional width of integument epithelial cells was significantly (13 %) greater in the acid-acclimated larvae compared to acid naïve larvae. Mucus secretion rates appeared to be significantly greater in both the gills and integument of acid-acclimated larvae (Table 3). Electron-dense mucus released from mucus secretory vesicles adhered to the apical surface of cells (Fig 4d). MSV density, total MSV cross-section area and the number of patent mucus vesicles were all much higher in acid acclimated larvae compared with acid naïve larvae. The total cross-sectional area of MSVs of acid acclimated larvae was over twice that of acid naïve larvae. The density of patent MSVs at the gill surface was over three times that of acid naïve larvae.

**Table 2.** Epithelial junctional morphology at the gill and integument surfaces in acid naïve and acid-acclimated larvae. Values shown are means ± s.e.m. N = 5 for all measures.

| Parameter | Treatment Group |  |
| --- | --- | --- |
|  | Acid naïve | Low pH-acclimated |
| *Branchial Epithelium* |  |  |
| Total junction length (μm) | 0.64 ± 0.03 | 0.78 ± 0.02* |
| Tight junction length (μm) | 0.37 ± 0.03 | 0.50 ± 0.04* |
| Minimum junction width (μm) | 0.010 ± 0.001 | 0.012 ± 0.001 |
| Maximum junction width (μm) | 0.028 ± 0.001 | 0.034 ± 0.002* |
| *Integument Epithelium* |  |  |
| Total junction length (μm) | 0.68 ± 0.03 | 0.82 ± 0.05* |
| Tight junction length (μm) | 0.33 ± 0.02 | 0.37 ± 0.04 |
| Minimum junction width (μm) | 0.011 ± 0.001 | 0.012 ± 0.001 |
| Maximum junction width (μm) | 0.031 ± 0.001 | 0.035 ± 0.001 |

\* indicates value was significantly different from that of acid naïve group ( $P < 0.05$ ).

**Table 3.** Branchial and epidermal vesicular mucus production / secretion in acid naïve and low pH-acclimated *Litoria cooloolensis* larvae. MSV = mucus secretory vesicle; CCA = cell cross-sectional area; TMSVA = total cross-sectional area of mucus secretory vesicles. Values shown are means ± s.e.m. N = 5 for all measures.

| Parameter | Treatment |  |
| --- | --- | --- |
|  | Acid naïve | Low pH acclimated |
| *Branchial Epithelium* |  |  |
| MSV density (MSVs per $\mu\text{m}^2$ CCA) | 0.073 ± 0.0017 | 0.119 ± 0.008* |
| TMSVA/CCA | 0.065 ± 0.009 | 0.181 ± 0.012* |
| Patent MSVs per $\mu\text{m}$ of apical cell membrane | 0.065 ± 0.009 | 0.233 ± 0.031* |
| *Integument Epithelium* |  |  |
| MSV density (MSVs per $\mu\text{m}^2$ CCA) | 0.097 ± 0.016 | 0.102 ± 0.012 |
| TMSVA/CCA | 0.049 ± 0.010 | 0.156 ± 0.025* |
| Patent MSVs per $\mu\text{m}$ apical cell membrane | 0.108 ± 0.018 | 0.308 ± 0.036* |

\* indicates value was significantly different from that of acid naïve group ( $P < 0.05$ ).

## Discussion

The tolerance of *L. cooloolensis* embryos and larvae to waters of low pH is unusual amongst amphibians, with acid waters pH 4 and less causing significant embryonic and larval mortality in most other species (see Freda, 1986; Rowe and Freda, 2000). The capacity of *L. cooloolensis* larvae to withstand chronic exposure to dilute waters as acidic as pH 3.5 is particularly noteworthy and on a par with fish species native to the extremely ion-poor and acidic Rio Negro (Gonzalez et al., 1997; Gonzalez and Dunson, 1989b; Gonzalez and Preest, 1999) and acidic lakes of Japan (Hirata et al., 2003). Compared to the closely related acid-sensitive sibling species, *Litoria fallax* (Meyer et al 2010), the capacity for survival at low pH in *L. cooloolensis* larvae may be a combination of both inherent species-specific traits and physiological plasticity.

Compared with larvae of the related, yet acid-sensitive species, *Litoria fallax* (Meyer et al., 2010), larvae of *L. cooloolensis* were substantially more tolerant of acute low pH exposure. In *L. fallax*, acute exposure of acid naïve larvae to pH 3.5 water resulted in 100% mortality within 24 h (Meyer et al. 2010) compared with 20% of *L. coolooensis* larvae acutely exposed to pH 3.5. Although exposure to pH 3.5 water did result in a substantial reduction in body Na^+^ content, surviving *L. cooloolensis* larvae only lost ∼30% of their total body Na^+^ in 24 h compared to the > 60% loss in moribund *L. fallax* larvae. As with previous studies (Freda and Dunson, 1984; Freda and Dunson, 1986a; Freda and Dunson, 1986b; McDonald et al., 1984; Meyer et al., 2010), those *L. cooloolensis* larvae that lost more than 50% of their body Na^+^ died. While the average body sodium content of larvae reared at pH 3.5 was lower than that of larvae maintained at pH 6.5, flux data indicate no net loss of body Na^+^ at this pH, with acid acclimated animals able to resist inhibition of Na^+^ uptake at pH 3.5 (unlike acid naïve larvae which suffered significant inhibition of Na^+^ uptake at pH 3.5).

In more acid-sensitive amphibians species like *Litoria fallax*, larvae exposed to pH 3.5 water experience not only inhibition of Na^+^ uptake but also increased Na^+^ efflux, resulting in a greater net loss of body Na^+^ (Meyer et al., 2010). In these species, the net loss of body Na^+^ can significantly disrupt ionic homeostasis causing larvae to die (see, e.g., Meyer et al., 2010). Survival of amphibian larvae at such low pHs therefore requires the regulation of both Na^+^ efflux and uptake rates, such that ionic homeostasis is maintained. Acute exposure of acid-sensitive amphibian larvae to low pH water exacerbates rates of Na^+^ efflux likely as a result of the disruption of epithelial paracellular junctions (Meyer et al., 2010). In previous studies, exacerbated Na^+^ loss at low pH has been attributed to the opening and shortening of epithelial tight junctions promoting the paracellular efflux of Na^+^ ions (Ferreira and Hill, 1982; Freda and Dunson, 1984; Freda et al., 1991; McDonald, 1983; Meyer et al., 2010). The comparatively low rates of Na^+^ efflux in *L. cooloolensis* larvae acutely and chronically exposed to pH 3.5 water suggest that the functional integrity of these junctions is unaffected by acid exposure (Ferreira and Hill, 1982; Freda and Dunson, 1984; Freda et al., 1991; McDonald, 1983; Meyer et al., 2010). In this case, resistance of *L. cooloolensis* larvae to increased Na^+^ efflux at low pH is not obviously related to tight junction depth or the length of junctional complexes between epithelial cells of the gills or integument. Resistance of *L. cooloolensis* larvae to increased efflux may, however, be linked to better apposition of gill epithelial cells, with junctional widths greater in acid-sensitive *L. fallax* larvae compared with *L. cooloolensis* larvae (Meyer et al., 2010). Differences between these species (in regards to their ability to resist increased Na^+^ efflux at low pH) could also be related to the composition of junctional proteins linking epithelial cells, with shifts in the composition of junctional proteins such as claudins and occludin having been shown to correlate with a decrease in Na^+^ efflux during low pH exposure in zebrafish (Kumai et al. 2011). Further work is required to determine if changes in these proteins also accompany acclimation/adaptation to low pH in *L. cooloolensis* larvae.

Protection of epithelial junction integrity during low pH exposure is well known to be influenced by external [Ca^2+^]. In the current study, water [Ca^2+^] significantly influenced Na^+^ efflux rates at low pH in both acid naïve and acid acclimated *L. cooloolensis* larvae, indicating that [Ca^2+^]_e_ is likely involved in maintaining junctional integrity. Provision of additional calcium offset some of the effect of pH on Na+ efflux rates suggesting that calcium handling (i.e. binding to junctional proteins, transcellular uptake) may be compromised at low pH. A similar protective effect of elevated [Ca^2+^]_e_ has been demonstrated for larval *Rana pipens* where Na^+^ efflux rates were suppressed almost completely (87%) following acute exposure to high Ca^2+^, low pH water (Freda and Dunson, 1984). Likewise, studies of acid tolerant fish have shown that in most cases, the presence of elevated [Ca^2+^]_e_ levels reduces Na^+^ efflux rates in waters of low pH, resulting in substantially reduced mortality rates (Glynn et al., 1992; Gonzalez and Dunson, 1989a; Gonzalez et al., 1998; Matsuo and Val, 2002; McWilliams, 1983; see also review by Walker et al., 1988), suggesting that tolerance of low pH environments is at least somewhat calcium-dependent. However, an understanding of the mechanistic basis by which [Ca^2+^]_e_ allows acid-tolerant species overcome the physical challenges imposed by low pH environs is lacking. The discovery of apical pH- and [Ca^2+^]_e_- sensitive Ca^2+^ channels (ECaC) in fish gill MR- and pavement-cells (Pinto et al., 2010; Qiu and Hogstrand, 2004; Shahsavarani et al., 2006) provides a potential molecular mechanism through which epithelial function might be both disrupted by low pH in acid sensitive species and improved in acid-tolerant species. Further work is needed to understand the mechanistic basis for improved resistance to Na^+^ efflux at low pH caused by [Ca^2+^]_e_ in amphibian larvae.

The protection of cell function and junctional integrity from the damaging effects of low pH water are central to tolerance of acidic waters. Relative to acid naïve larvae, high levels of mucus production and secretion were seen in acid-acclimated larvae. Mucus has several well established roles including the development of an unstirred layer and facilitation of ion/water transport (Handy et al., 1989; Shephard, 1981; Shephard, 1982), as an immunological barrier (Swidsinski et al., 2007) and in the control of gas exchange across epithelia (Wright et al., 1989). Mucus production is often stimulated by noxious substances such as low pH (Bohmer and Rahmann, 1990; Leino and Mccormick, 1984; Linnenbach et al., 1987; Walker et al., 1988), where its primary function is likely increasing the distance between the cell surface and the environment (Henriksnas et al., 2006; Phillipson et al., 2008) and neutralising hydrogen ions before they reach the apical surface of epithelial cells (Holma, 1985; Phillipson et al., 2008; Stith, 1984a; Stith, 1984b). Other studies have demonstrated an important role for branchial mucus in concentrating electrolytes within an unstirred layer, thereby preventing ion leak from epithelia (Handy et al., 1989) and also promoting Na^+^ uptake (Stith, 1984a; Stith, 1984b). The diffusion of low-charge ions, like monovalent electrolytes, through the mucosal unstirred layer is many cases similar to that through normal saline indicating that mucus is not a major obstruction to the effective uptake of these ions (Handy and Maunder, 2010). However, there is some evidence to suggest that the mucus layer may increase the oxygen diffusion distance thereby limiting the capacity of tissues to effectively uptake oxygen (Saldena et al., 2000). Whether this translates into an increased cost of living at low pH in terms of growth rates or energy use, remains to be determined for acid water specialists like *L. cooloolensis* larvae.

Tolerance of low pH environments and the maintenance of ionic homeostasis requires that both Na^+^ loss and Na^+^ uptake rates be controlled. Unlike *L. fallax* larvae, acute exposure to low pH water did not result in significant inhibition of Na^+^ uptake in *L. cooloolensis* larvae, and indeed in *L. cooloolensis* larvae reared at pH 3.5, Na^+^ uptake rates were indistinguishable from those of acid naïve larvae at pH 6.5. These data suggest that the capacity to limit the acute and chronic impacts of low pH on ion uptake mechanisms in *L. cooloolensis* larvae contributed to their tolerance of acid water. Although the precise mechanism is not well established, based on morphological studies of transport epithelia in salamander larvae, and on frog skin (Koefoedjohnsen and Ussing, 1958; Uchiyama et al., 2011a; Uchiyama et al., 2011b; Ussing and Zerahn, 1951), the bulk of extra-digestive Na^+^ uptake probably occurs through membrane-bound apical epithelial Na^+^ channels (ENaC) at the gill surface, whose activity is coupled to the Na^+^ gradient generated by the activity of basolateral Na^+^K^+^ATPase pumps (Butterworth, 2010). In other amphibian tissues, a large decrease in apical pH (< pH 4) can cause a significant reduction in ENaC-mediated Na^+^ transport (Harvey et al., 1988; Palmer, 1985; Ussing and Zerahn, 1951; Zeiske et al., 1999). A reduction in apical pH is also likely to lower intracellular pH as a consequence of H^+^ permeation, especially if cellular H^+^ extrusion mechanisms are unable to keep up with H^+^ infiltration rates. *In vitro* models of ENaC activity indicate that Na^+^ transport activity is altered primarily by changing the channel open probability (P_o_) or by changing the abundance of ENaC pumps (N), both of which decrease with decreasing intracellular pH (Palmer and Frindt, 1987; Zeiske et al., 1999). Although an acute reduction in pH likely lowered ENaC-mediated Na^+^ uptake in *L. cooloolensis*, restoration of Na^+^ uptake capacity with prolonged exposure to low pH may permit long-term habitation of low pH environments by *L. cooloolensis* larvae.

Maintenance of Na^+^ uptake in acid-acclimated *L. cooloolensis* larvae appears to have been accomplished by an increase in maximum Na^+^ transport capacity, due most likely to an increase in the number of open Na^+^ channels in the cell (as opposed to an increase in the affinity of channels for Na^+^ which would have lowered K_m_). As with *L. cooloolensis* larvae, the ability of many acidophilic fish to maintain Na^+^ uptake at low pH also appears attributable to increased capacity for Na^+^ transport, in addition to a high transporter affinity for Na^+^ (low K_m)_ for Na^+^ (Gonzalez and Preest, 1999; Gonzalez and Wilson, 2001). The lower K_m_ of these acid tolerant fish species is considered a fixed trait reflecting evolutionary adaptation to salt-depleting environs (Ultsch et al., 1999). It is likely that in acid-acclimated *L. cooloolensis* larvae, an increase in the abundance of Na^+^ transporting pumps is required to maintain homeostasis against the persistent leak of Na^+^ from the body that occurs in low pH water. In addition, H^+^ extrusion mechanisms may be upregulated to better control intracellular pH, as has been observed in zebrafish (Hirata et al., 2003; Horng et al., 2009), which may improve the transport efficiency of Na^+^ channels operating in low pH environments. Indeed, in the absence of intracellular acidification, exposure of the extracellular components of ENaC to low pH bathing solution can improve ENaC P_o_ and, therefore, Na^+^ uptake capacity (Wichmann et al., 2019). The capacity to acclimate ion uptake rates to low pH with prolonged exposure suggests that this system is highly plastic which allows animals to match ion uptake requirement to the prevailing environment. Our findings are also consistent with several studies showing that acid-induced disturbances to Na^+^ transport mechanisms can be recovered over time in some fish species (Gonzalez and Dunson, 1989a; Gonzalez and Dunson, 1989c). The capacity to manage intracellular acidification could be a major factor underpinning acclimation and adaptation to low pH environments in *L. cooloolensis* larvae, however further work to describe the molecular basis for Na^+^ and H^+^ ion transport in amphibian larvae is required.

Based on the results of this study, the tolerance of *L. cooloolensis* larvae to low pH may be attributed to a combination of physiological and morphological factors promoting resistance to acid damage and disruption of ionic homeostasis in soft acid waters. This includes a high capacity for branchial Na^+^ uptake, tighter apposition of epithelial cells of the gills and integument, and greater mucus production and secretion at the gills and body surface. These attributes appear to enable *L. cooloolensis* larvae to tolerate acid stress better than larvae of most other amphibian species. Since *L. cooloolensis* breeds naturally in acidic soft-water lakes, these traits are likely to reflect adaptation to soft, low pH environments. Although many questions remain, an understanding of the molecular identity and function of epithelial ion transport systems in *L. cooloolensis* and other similarly acid-tolerant animals is necessary to determine the precise nature of the mechanisms responsible for adaptation to low pH water.

